# Understanding the killing mechanism of action by virus-infected yeasts

**DOI:** 10.1101/398099

**Authors:** Sean Sheppard, Duygu Dikicioglu

## Abstract

Killer yeasts are microorganisms, which can produce and secrete proteinaceous toxins, a characteristic gained *via* infection by a virus. These toxins are able to kill sensitive cells of the same or a related species. From a biotechnological perspective, killer yeasts have been considered as beneficial due to their antifungal/antimicrobial activity, but also regarded as problematic for large-scale fermentation processes, whereby those yeasts would kill species off starter cultures and lead to stuck fermentations. Here, we propose a mechanistic model of the toxin-binding kinetics pertaining to the killer population coupled with the toxin-induced death kinetics of the sensitive population to study toxic action *in silico*. Our deterministic model explains how killer *Saccharomyces cerevisiae* cells distress and consequently kill the sensitive members of the species, accounting for the K1, K2 and K28 toxin mode of action at high or low concentrations. The dynamic model captured the transient toxic activity starting from the introduction of killer cells into the culture at the time of inoculation through to induced cell death, and allowed us to gain novel insight on these mechanisms. The kinetics of K1/K2 activity *via* its primary pathway of toxicity was 5.5 times faster than its activity at low concentration inducing the apoptotic pathway in sensitive cells. Conversely, we showed that the primary pathway for K28 was approximately 3 times slower than its equivalent apoptotic pathway, indicating the particular relevance of K28 in biotechnological applications where the toxin concentration is rarely above those limits to trigger the primary pathway of killer activity.

## 1. Introduction

Killer yeasts are eukaryotic, single-celled fungi, which produce and secrete toxic proteins that are lethal to sensitive cells; the phenomenon was first characterised in 1963[1]. Several yeast species are recognised as possessing the killer characteristic with the most extensively studied one being *Saccharomyces (S.) cerevisiae* [2]. The potential of killer yeasts due to their antimicrobial activity has been explored widely, especially within the context of applications in the food industry [3]. Although being acknowledged as a promising premise for a number of biotechnological applications, killer activity of yeasts is also deemed undesirable on other platforms.

These infected cells are often detected in wine fermentation processes whereby killer yeasts contaminate starter cultures, killing the microbiological fermenting agents. Ratios as low as one killer yeast for every hundred sensitive yeasts was reported to eliminate the starter culture population within 24 hours [4–6]. Stuck fermentations, which are characterised by high concentrations of acetaldehyde and lactic acid, are often a consequence, and is typical of a very distasteful wine product [7]. Wineries can incur substantial financial losses due to stuck fermentations, and therefore, killer yeasts are considered as an important concern compromising success in commercial wine processing [8]. Early research on wine microbial communities focused on devising control strategies to manage killer yeasts in wine fermentation in order to reduce the chance of spoilage [2]. More recently, the possibility of commercialising killer yeasts in wine fermentation specifically due to their antifungal activity was also investigated [9]. Within this remit, genetic modification techniques were explored as potential tools for constructing wine yeast strains, which are resistant to, and/or possess the killer factor [10]. Starter cultures, into which these strains would successfully be integrated, could assist the control and prevention of contaminating fungal species *via* the production of toxins, making wine production more reliable than before, and reducing the costs associated with stuck fermentations substantially [11,12].

Killer yeast systems are classified on the basis of the molecular characteristics of their toxins, the variations in the encoding genetic determinants, the presence or the lack of cross-immunity, and their killing profiles [13]. *S*. *cerevisiae* is by far the most studied yeast species with regards to acquired toxic activity, with four distinct toxin proteins characterised (K1, K2, K28 and Klus), thus it contributes vastly to our understanding of killer yeasts and their infecting viruses. The virus, L-A, is an icosahedral double stranded (ds) RNA virus of the yeast *S. cerevisiae* with a single 4.6 kb genomic segment that encodes its major coat protein, Gag (76 kDa) and a Gag-Pol fusion protein (180 kDa) formed by a -1 ribosomal frameshift, which encode an RNA-dependent RNA polymerase. A number of satellite dsRNAs, called M dsRNAs encoding a secreted protein toxin and immunity to that toxin are hosted in separate viral particles, whose replication and encapsidation are supported by L-A. Each toxin is encoded by a single open reading frame, and is synthesised as a polypeptide preprotoxin. The preprotoxin comprises a hydrophobic amino terminus representative of secretion and is modified *via* the endoplasmic reticulum and the Golgi apparatus of the host for activation of the toxin prior to its secretion [8,17– 17].

Only those yeasts, which possess both the M dsRNAs and L-A, are able to produce effective toxins. Although the transmission of L-A and M from cell to cell occurs exclusively during the mating process but not *via* natural release from the cell or entry by another mechanism, it is the high frequency of yeast mating, which ensures the wide distribution of these viruses in natural isolates. Moreover, the structural and the functional similarities of these viruses to dsRNA viruses of mammalian systems evoke further interest in their study [14,17,18].

K1 is comprised of α-(9.5 kDa) and β-subunits (9.0 kDa) [13]. The β-subunit is responsible for receptor binding to sensitive cells, and the α-subunit induces the lethal effect. K2 structurally resembles K1 with subunits that are only marginally larger than those of K1[19], despite substantial differences in the sequences of their dsRNA, their molecular weights, isoelectric points and optimum pH [20]. The mechanisms through which these toxin proteins kill susceptible cells display some level of variance, although striking similarities also exist. K2 toxin, less extensively characterised than K1, was accepted to have a similar mechanism of action to K1, apart from some differences that occur at the plasma membrane and cell wall level. The patterns of processing of both killer proteins by the sensitive cells were also reported to be similar. Despite the similarities of their toxins, K1 and K2 killer strains were reported to be able to kill each other even though they are immune to their own toxin [21]. K28 is also secreted as a heterodimer of α-(10.5 kDa) and β-subunits (11 kDa) [22]. The mechanism of K28-induced lethality was reported to be significantly different than that of K1-induced lethality [23]. Klus was recently isolated as a novel toxin produced by *S*. *cerevisiae* [24], and its toxic action is still understood very little.

Cells that are susceptible to these toxins were identified to possess two different types of sites or receptors that bind the killer toxin with different affinities [25]. The first step of K1/K2 binding was reported as the low affinity, high velocity, energy independent adsorption of the killer toxin on β-1,6-D-glucans embedded in the cell wall [26,27]. Once bound, the toxin had a high affinity, low velocity, energy-dependent interaction with Kre1p receptors; the glycosyl-phosphatidylinositol-linked glycoproteins located on the cell membrane [28,29]. Subsequently, the α-subunit of the toxin was shown to trigger the formation of voltage-independent cation transmembrane channels, which would then cause the leakage of H^+^ and K^+^ ions, followed by cell death [30]. In contrast to K1 or K2, K28 was reported to initially bind to α-1,3-linked mannose residues of a 185-kDa cell wall mannoprotein [31]. The toxin would then interact with the Erd2p receptor triggering toxin uptake into the cytosol by receptor-mediated endocytosis [32]. The β-subunit of K28 would be ubiquitinated and proteasomally degraded following the toxin’s uptake and retrograde transport through the Golgi and endoplasmic reticulum to the cytosol, while the α-subunit cleaved from its β-subunit and migrated into the sensitive cell’s nucleus [33]. There the lethal subunit would arrest cell cycle at the G1/S boundary, preventing the separation of the daughter and the mother cells, effectively killing both cells [33,34].

The primary mechanisms of action for these toxins are in place at high concentrations of toxin availability. A secondary mechanism was described relatively recently, and was shown to be in effect upon medium to low-concentration exposure to toxins. The toxin proteins were shown to induce programmed cell death at concentrations that were not sufficient to trigger the primary action pathways [11,35–38]. The affected sensitive cells were shown to enter an apoptotic state; exhibiting markers such as DNA fragmentation, chromatin condensation and phosphatidylserine externalisation [37]. The activation of the yeast metacaspase 1 (Yca1p) *via* toxins would eventually yield to the release of reactive oxygen species, triggering a cascade of events consigning the cell to death *via* apoptosis [39,40]. This response appears to be universally induced at low concentrations, irrespective of the nature of the toxin protein in question. Toxic action *via* programmed cell death has been proposed as the predominant mechanism by which killer yeasts kill sensitive species in natural environments where they are found in much lower concentrations [41].

Despite the implications on the importance of the secondary mechanism of toxic action in killing potential, questions such as the threshold at which the switch between the primary and the secondary mechanisms occur, or how the toxin binding kinetics were related to the toxin-induced death kinetics, still remain open. In this work, we built a deterministic model that could simulate both the toxic activity of killer cells, and the toxin-induced damage on the sensitive population upon exposure. The model equipped us with a novel platform to (i) study the toxic effects of killer yeasts on sensitive populations, (ii) predict how the system responds to varying levels of toxin, and (iii) develop strategies to control and contain killer population and maintain the survival of the sensitive population. We selected both the killer and the susceptible population to be comprised of *S*. *cerevisiae*, and compiled the extensive qualitative and quantitative data available in the literature on the kinetic action of the K1, K2 and K28 toxins at high and low concentrations. We implemented a model structure and verified the models’ predictive power on independent datasets; they were able to accurately match empirical data. We then employed the models to gain in-depth novel insight into how the mechanisms for toxic activity changed in response to the extent of toxin exposure. The constructed models are publicly available via EBI’s BioModels Database[42] *via* access no’s: MODEL1804230001 MODEL1804230002 for K1/2 and K28, respectively.

## 2. Materials and Methods

### 2.1. Model structure and modelling platform

The model described the killer toxin activity, which comprised of the binding phase of the toxin to the cell wall and membrane, and the sensitive cell activity, which comprised of toxin-induced death of the sensitive fraction of the population. These activities explaining the mechanism were represented by chemical reactions between molecular species and the dynamics were governed by rate laws associated with each reaction. The model was built using ordinary differential equations. The model was built and simulated in the open-source software COPASI 4.22 (Build 170) [43]. The timescale of the analysis was measured in minutes as it allowed the observation of both the rapid toxin binding, and the relatively slow toxin-induced death in the same frame. The concentration of both the toxins and cells were represented as the number of molecules per ml (molecules/ml).

### 2.2. Experimental data

Experimental data was acquired from relevant publications. Whenever data was only presented in the form of Figures in their respective publications, the open-source Java platform Plot Digitizer 2.6.8 (http://plotdigitizer.sourceforge.net/) was employed to get an accurate estimate of the raw data.

The specific masses of the toxins were acquired from [13] as 19000 Da for K1, 21500 Da for K2, and 21500 Da for K28; consequently, 1 pg of toxin equated to 3.125×10^7^, 2.8×10^7^, and 2.8×10^7^ molecules of K1, K2, and K28, respectively. Data from *Pichia membranifaciens* PMKT and PMKT2 toxins, which were reported to be analogous to K1 and K28 were employed to infer kinetic information and to complement data on *S*. *cerevisiae* toxins when no direct data was available [11,44,45]. Experiments from which data was collected from were carried out under the ideal conditions for each toxin and so the same conditions are assumed to exist for the model. There was an overlap of the optimal temperature and pH for all three toxins, and all data employed in the study were reported at the same conditions of pH = 4.7 and T = 22 °C [30,46,47]. The viable cell population was considered to be at stationary phase where the natural death and growth rate of the population are equal, so population growth was excluded from the model structure, since data was available on such populations at dynamic equilibrium, in order to detect the extent of death caused by the toxins.

### 2.3. Representation of the toxin binding phase

The main underlying assumption of the binding model was that it assumed a well-mixed population of sensitive cells. Toxin proteins were also assumed to be well-mixed within sensitive cells, so there was no heterogeneity to how toxins would bind to individual cells, but instead would accumulate simultaneously across all cells in the population. 16,800,000 molecules of K1 were calculated to saturate the cell wall receptors of a single *S*. *cerevisiae* cell from the data provided in [25]. The cell membrane Kre1p receptor was reported to be saturated at a concentration 50-fold lower than that needed to saturate the cell wall receptor [26] yielding a saturation constant of 336,000 molecules. The kinetics of K1 binding to the cell membrane was reported to be 6.333 times slower than binding to the cell wall [25]. Kinetic data for cell wall and cell membrane binding were available for K1 and K2 (Table 1), but not for K28. However, the number of cell wall mannoproteins was reported to be similar to that of β-1,6-D-glucans [48] and K28 has a similar mass to K1 and K2. The saturation constants, cell wall and cell membrane binding rates were assumed as identical, since no data was available to contradict this assumption. Furthermore, the binding rate of PMKT2 (K28 equivalent toxin) to α-1,3-linked mannose residues was reported to have similar kinetics to K1 [11], providing further support to the assumption made.

**Table 1.**
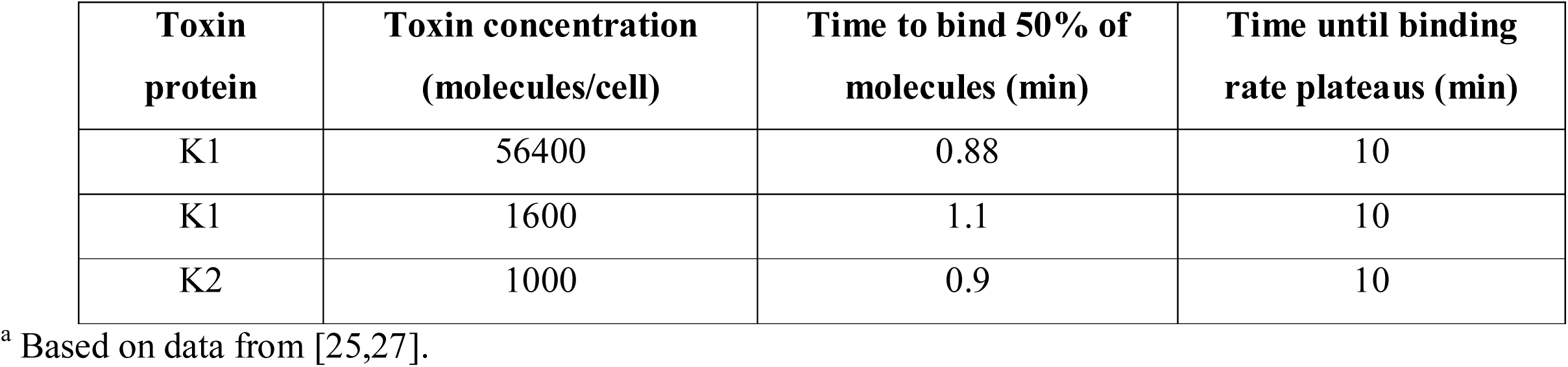
Kinetics of Toxin Cell Wall Binding^a^

| Toxin protein | Toxin concentration (molecules/cell) | Time to bind 50% of molecules (min) | Time until binding rate plateaus (min) |
| --- | --- | --- | --- |
| K1 | 56400 | 0.88 | 10 |
| K1 | 1600 | 1.1 | 10 |
| K2 | 1000 | 0.9 | 10 |

<sup>a</sup> Based on data from [25,27].

### 2.4. Representation of toxin-induced cell death

An individual *S*. *cerevisiae* cell of average sensitivity was reported to be killed by at least 28000 K1or K28 molecules [26,49], which was selected as the threshold value for the activation of the primary pathway. Below this concentration (28000 molecules/cell), the model activated the secondary apoptotic pathway, as was reported earlier [37]. Toxin-induced cell death was reported to be ineffective at toxin concentration of 852 K1 molecules/cell and 1080 K28 molecules/cell [37]. The threshold at which the cells displayed the first signs of apoptotic cell death upon exposure to toxin was adopted from the investigation of PMKT2 activity, where, at this threshold, approximately 35% of the cells showed signs of entering apoptosis after 90 minutes upon toxin exposure, with the ratio rising to 95% after 180 minutes [11]. A similarity was assumed between the K1/2- and K28-induced cell death kinetics due to the now well-acknowledged similarities between the mechanisms of toxic action to induce apoptotic cell death. Kinetic data for toxin-induced death was available for both K1 and K28, but only at certain concentration levels (Table 2).

**Table 2.** Kinetics of Toxin-Induced Cell Death^a^

| Pathway | Toxin protein | Toxin concentration<br>(molecules/cell) | Time to kill 50% of<br>population (min) | Time to kill 98% of<br>population (min) |
| --- | --- | --- | --- | --- |
| Primary | K1 | 75000 | 7 | 40 |
| Apoptotic | K1 | 2600 | 40 | 220 |
| Primary | K28 | $9.63 \times 10^9$ | 100 | 560 |
| Primary | K28 | $3.75 \times 10^6$ | 90 | 560 |
| Apoptotic | K28 | 2800 | 40 | 230 |
| Primary | PMKT2 | 28000 | 120 | 570 |

<sup>a</sup> Based on data from [11,34,37,50].

## 3. Results

A single model structure was employed to describe K1/K2 kinetics and K28 kinetics, with relevant constants to account for the difference in rates of toxin binding and toxin-induced cell death (see Table 3 for constants). A modular system of toxin binding and killing activities was implemented, and the model complexity was increased in a stepwise manner such that the final working model either made use of (during re-construction), or explained (during benchmarking and validation) virtually all of the empirical data available to date.

**Table 3.** Parameters of the Rate Equations used as the Basis of the Model^a^

| Biological Process | Parameter | Value | Units |
| --- | --- | --- | --- |
| Cell wall binding | $V_{max_{CW}}$ | 1.23 | *molecules/L · min* |
| Cell wall binding | $K_{CW}$ | $9.07 \times 10^9$ | *molecules/L* |
| Cell membrane binding | $V_{max_{CM}}$ | 0.205 | *molecules/L · min* |
| Cell membrane binding | $K_{CM}$ | $9.07 \times 10^9$ | *molecules/L* |
| Toxin-induced cell death *via* primary pathway (K1 & K28) | $LU_{Primary}$ | 28000 | LU<sup>b</sup> |
| Toxin-induced cell death *via* apoptotic pathway (K1) | $LU_{Apoptotic(K1)}$ | 825 | LU<sup>b</sup> |
| Toxin-induced cell death *via* apoptotic pathway (K28) | $LU_{Apoptotic(K28)}$ | 1080 | LU<sup>b</sup> |
| K1-induced death | $V_{max_{TID(K1)}}$ | $5.0 \times 10^{-7}$ | *molecules/L · min* |
| K28-induced death | $V_{max_{TID(K28)}}$ | $2.5 \times 10^{-13}$ | *molecules/L · min* |
| Adjusted K1-induced death | $V_{max_{ATID(K1)}}$ | $2.5 \times 10^{-9}$ | *molecules/L · min* |
| K1-induced apoptosis | $V_{max_{TIA(K1)}}$ | $4.0 \times 10^{-9}$ | *molecules/L · min* |
| K28-induced apoptosis | $V_{max_{TIA(K28)}}$ | $5.0 \times 10^{-9}$ | *molecules/L · min* |
| K1-triggered apoptotic death | $V_{max_{AD(K1)}}$ | 0.30 | *molecules/L · min* |
| K28-triggered apoptotic death | $V_{max_{AD(K28)}}$ | 0.30 | *molecules/L · min* |

<sup>a</sup> If the value of a parameter was obtained by parameter scanning, the median value is displayed above.
<sup>b</sup> LU: lethal units

### 3.1. Molecular crowding around cell wall receptors impedes binding at high toxin abundance

A common binding model proposing similar binding mechanisms for K1 and K28 was constructed, as suggested by empirical data reported in previous studies, and the rate of binding of the toxin on the yeast cell wall was captured by parameter estimation from [25]. Binding of the toxin molecules on the cell wall receptors and on the cell membrane was modelled separately. The simplest preliminary model for binding was constructed only to account for the phenomenon occurring on the cell wall. Michaelis-Menten kinetics, often employed to quantify the degree of interaction between a ligand and a receptor population [51], was selected for this model (eq. 1), but it failed to capture the initial dynamics of cell wall binding, overestimating the amount of toxin bound (Fig, 1). This simple model did not account for any crowding of toxin molecules at the receptor binding sites, and therefore allowed binding without any constraints as long as unbound toxins were available. Consequently, the possibility of any potential competition at the receptor binding sites, which could slow binding down by restricting access to the receptors despite their availability was overruled in this initial model.

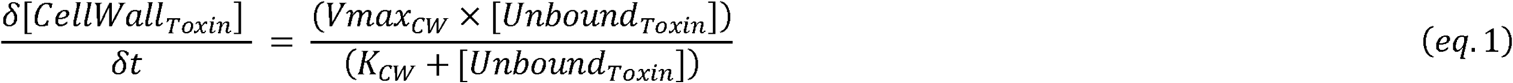

In order to account for this inefficiency in binding, the rate law was modified to (eq. 2), improving the simulation of the binding characteristics observed during the first two minutes following the exposure of sensitive cells to toxin molecules (Fig. 1a). This improvement suggested that a potential interaction between the unbound toxin molecules and those that were already bound to the cell wall receptors could indeed hinder binding within the initial minutes upon exposure.

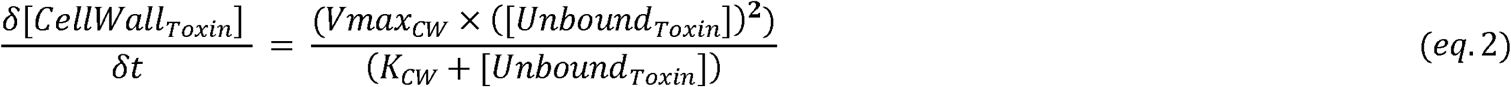

**Fig. 1.**
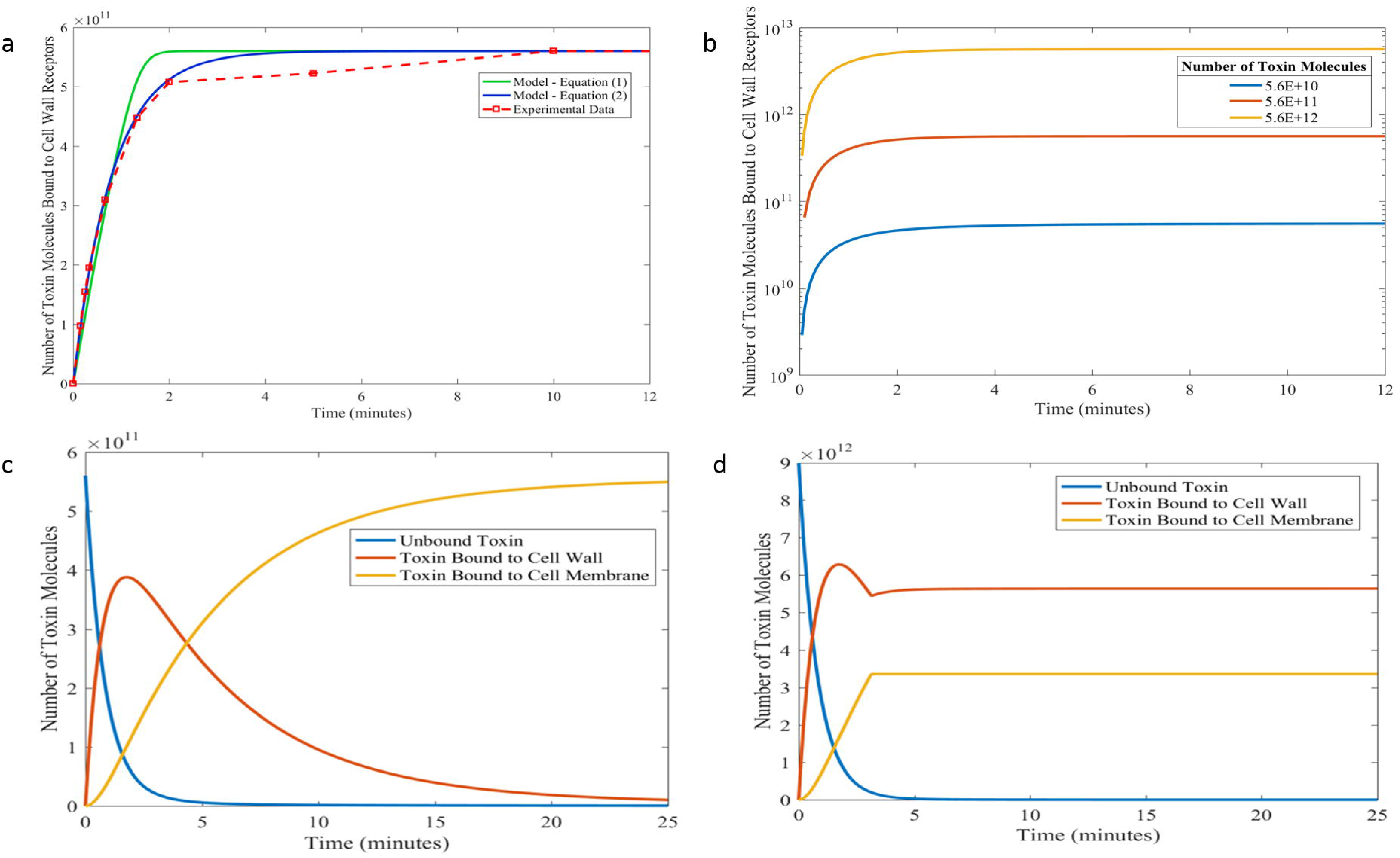
Binding kinetics of toxin molecules. (a) Kinetic models of cell wall binding of toxins employing (eq. 1) shown in green and (eq. 2) shown in blue. Experimental data shown in red (data points connected with a dashed line) was taken from [25]. Initial population of unbound toxins was fixed at 5.6×10^11^ molecules. (b) Simulations using the kinetic model for cell wall binding of toxin molecules (eq. 3) starting from different initial concentrations of unbound toxins. The initial unbound toxin concentration was explored across a range of 2 orders of magnitude. (c) Predictive model simulations explaining the kinetics of K1, K2 and K28 binding to the cell wall and the cell membrane with unsaturated receptors available on both surfaces. The initial toxin availability was 5.6×10^4^molecules/cell. (d) Predictive model simulations explaining the kinetics of K1, K2 and K28 binding to the cell wall and the cell membrane with receptors on both surfaces reaching full saturation. The initial toxin availability was 9×10^5^ molecules/cell. Note that the total number of toxin molecules for 1×10^7^ susceptible cells are displayed in (a-d).

The saturation of the cell surface receptors was described by a saturation constant such that if the concentration of bound toxin was higher than the value of that constant, no further binding would take place (eq. 3). This allowed the representation of a step-wise binding process that was dependent on the availability of toxin molecules per each cell.

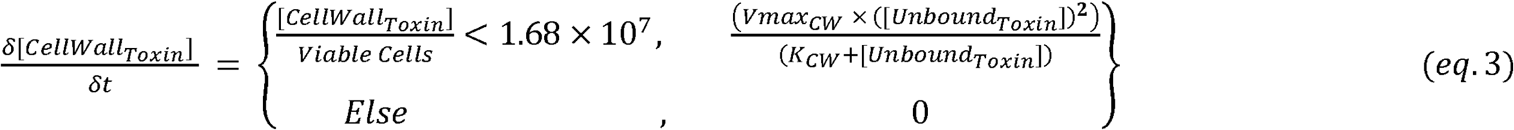

This model of cell wall binding kinetics was simulated for an arbitrarily selected population of 1×10^7^ susceptible cells that were exposed to varying initial concentrations of unbound toxins in order to investigate the effect of saturation on cell wall binding kinetics. The initial concentration of unbound toxins varied in a 100-fold range. The analysis showed that as long as the saturation of the receptors on the cell wall was avoided, the binding kinetics remained constant (Fig, 1b). Similar binding rates were reported in experiments where the initial unbound toxin concentration was varied by more than 50-fold, providing further support for these findings [25,27].

The cell wall binding kinetics for toxin molecules was extended also to describe cell membrane binding kinetics. Although the membrane binding mechanism was assumed to be similar to the mechanism for binding the cell wall, maximum specific rate of binding, *Vmax*_*CM*_, was reported to be 6.333 times slower [25]. The saturation of the cell membrane surface by the toxins was also described by a conditionality, and the final model for binding accounted for the unbound toxin molecules to initially bind to the cell wall receptors (represented in eq. 4), and then to be translocated to the cell membrane (represented in eq. 5).

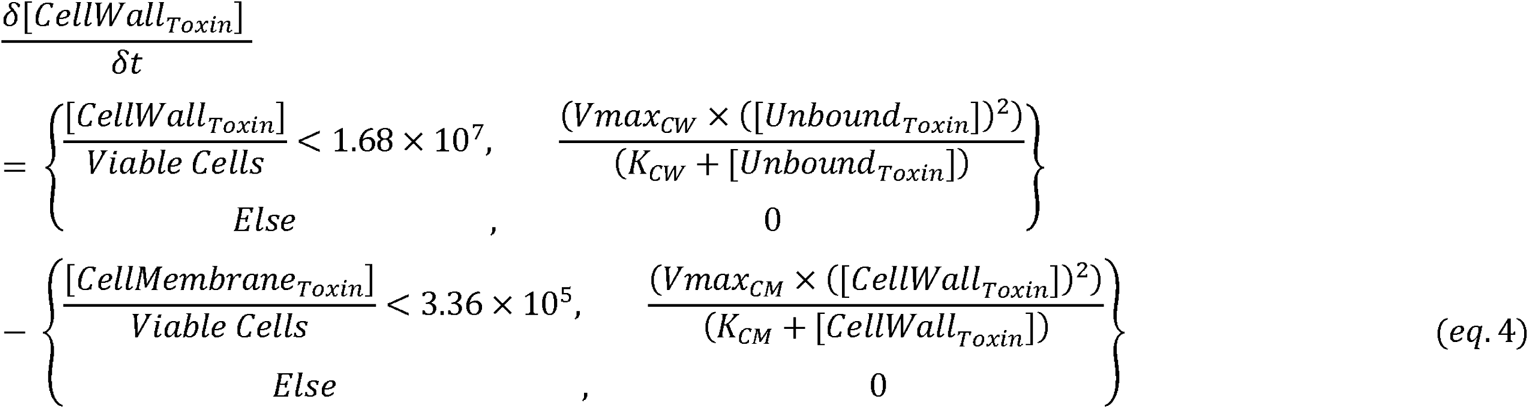

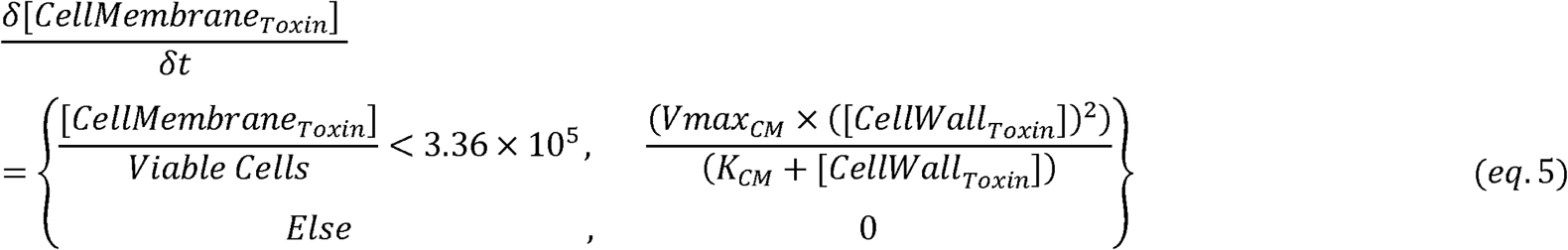

The model predicted that the extent of toxin binding to the cell wall rapidly increased and reached a maximum of 56,000 molecules/cell 103 seconds after the initial exposure to the toxin proteins. This bound toxin population was observed to shrink later, as toxin molecules slowly migrated towards the surface of the cell membrane. The toxin migration to the cell membrane was observed slow down approximately after 20 minutes leading towards the saturation of the membrane surface (Fig. 1c). The membrane receptor population was shown to fully saturate at a toxin concentration of 336,000 molecules/cell or higher, and no further toxin molecule movement was observed from the cell wall to the membrane, and two distinct toxin populations bound to different compartments of the cell were achieved (Fig. 1d).

### 3.2. Integrating toxin-induced death dynamics and toxin binding dynamics *via* the utilisation of lethality units

The model representing the kinetics of toxin binding to the cell wall and the cell membrane was further extended to represent the dynamics of cell death in a sensitive population exposed to toxins released by the killer population. A functional link was established between the toxin molecules and the cells that were doomed to die upon exposure to these molecules in order to facilitate the reconstruction of an integrated model. A new measure called the lethal unit (LU) was introduced into the model in order to represent a cluster of toxin molecules, which would be sufficient to induce toxin-associated death of a single yeast cell. The two different routes of toxin-induced death; via the primary pathway or via the apoptotic pathway required differently sized clusters of toxin molecules to be defined into a single LU. Therefore, the number of toxin molecules initially available per cell was used by the model to determine the pathway through which the toxins exerted their lethal effect. If the initial concentration was above 28000 toxin molecules/cell the model adopted the primary pathway of toxin activity to induce cell-death by converting every 28000 molecules into a single LU. Otherwise, 852 or 1080 toxin molecules leading the sensitive cell towards apoptosis were employed to represent a single LU for K1 and K28, respectively (eq. 6).

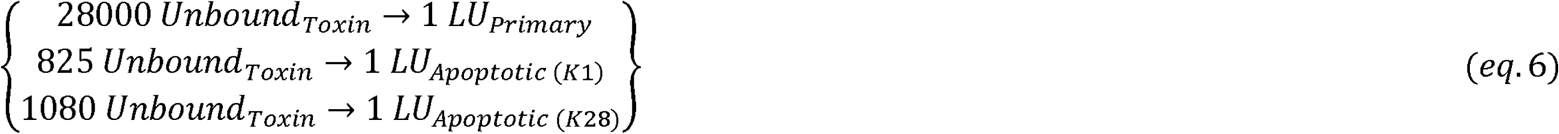

The efficacy of the toxins on the cell population was described by mass action kinetics. The cell death kinetics via the apoptotic pathway and the primary pathway of toxin activity were thus compared for both K1 and K28. Our model simple mass action kinetic analysis showed that K1 introduced at a concentration of 75,000 molecules/cell, killed off 50% of the sensitive population 7 minutes. In 40 minutes, 98% of the whole population was dead (Fig. 2a). The apoptosis-inducing toxin lethality acted considerably slower on the sensitive cells than the primary route of toxic activity. The presence of K1 at a concentration of 2600 molecules/cell could only kill 50% of the population after 40 minutes post initial toxin exposure, whilst it took as long as 220 minutes (> 3.5 hours) for 98% of the cells constituting the population to become inviable (Fig. 2b).

**Fig. 2.**
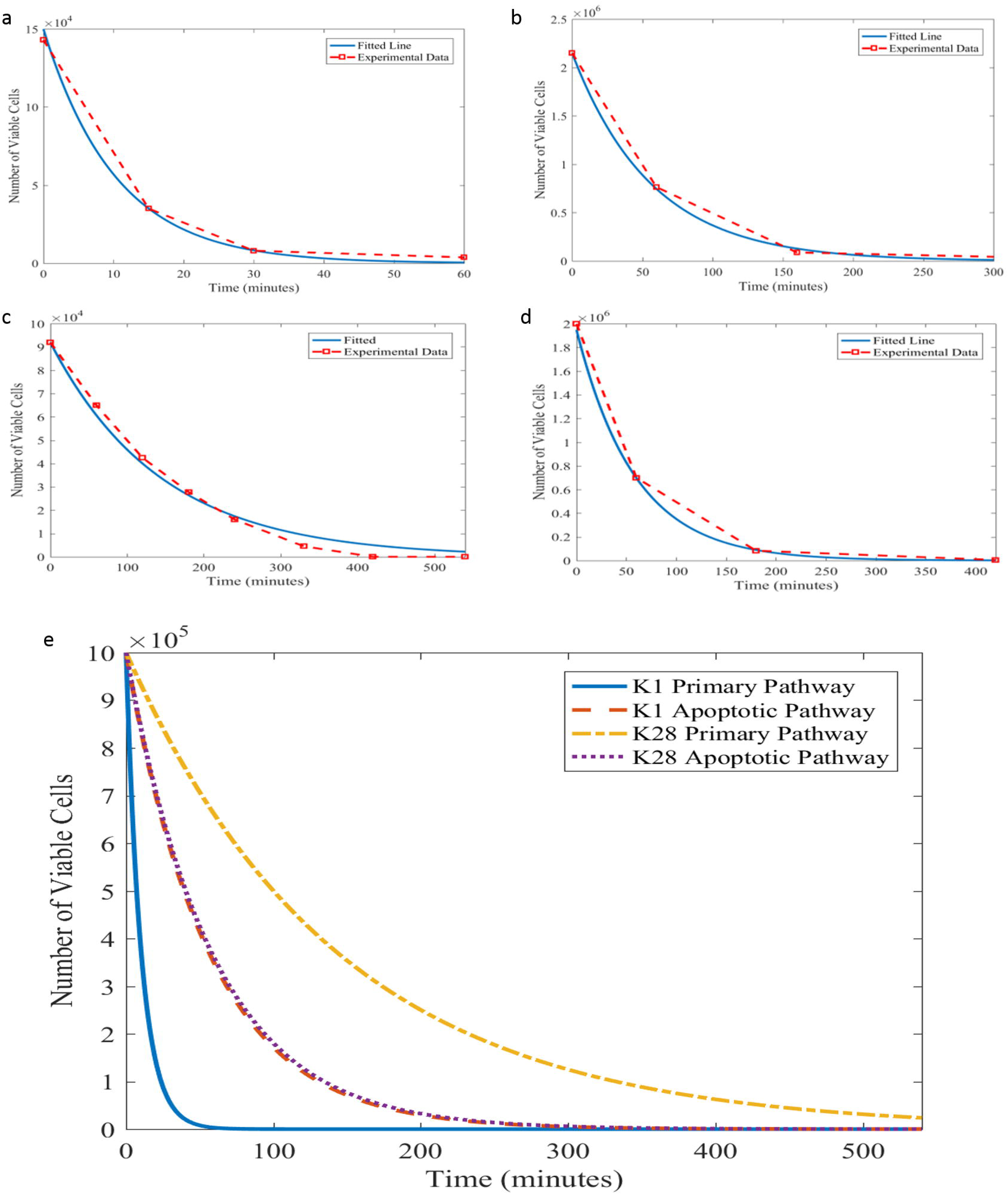
Simulation of toxin induced cell death. (a-b) Kinetic models of toxin-induced cell death by K1 for the primary pathway of toxin activity (a) and the apoptosis-inducing pathway of toxin activity (b). Experimental data shown in red (data points connected with a dashed line) was taken from [37]. Parameter estimation was employed to fit the data. Toxin concentration was 75000 molecules/cell in (a) and 2600 molecules/cell in (b). (c-d) Kinetic models of toxin-induced cell death by K28 for the primary pathway of toxin activity (c) and the apoptosis-inducing pathway of toxin activity (d). Experimental data shown in red (data points connected with a dashed line) was taken from [37,50]. Parameter estimation was employed to fit the data. Toxin concentration was 9.63×10^?^ molecules/cell in (c) and 2800 molecules/cell in (d). (e) Comparison of the Kinetics of the Toxin-Induced Cell Death Pathways for K1 and K28 are provided with the initial size of the viable population normalized to 150,000 for all four pathways.

K28 had a substantially slower primary effect to induce cell death than K1 did. The data showed that nearly 100 minutes was needed to reduce the viable cell population by 50% even at a toxin concentration as high as 9.63×10^9^ molecules/cell. More than 560 minutes was required for 98% of the cell population to become inviable (Fig. 2c). In contrast, the apoptotic pathway for K28 was nearly as fast as that for K1 to induce cell death, and in, fact, the pertaining kinetics were even faster than those for its primary pathway. The availability of 2800 K28 molecules/cell was sufficient to kill 50% of the population within 40 minutes, and 98% of the population was inviable after 230 minutes post exposure to toxin (Fig.2d).

The rates of toxin induced death displayed a nearly 16-fold difference across different toxins available at different concentrations. The apoptotic pathways of K1 and K28 were very similar, inducing death in 1.763% and 1.713% of the population per minute, respectively. This similarity, despite the substantial differences in the kinetic activity shown by their primary pathways, supported the hypothesis that toxin-induced apoptotic cell death could indeed be a universal mechanism of toxin activity (Fig. 2e). The primary pathway of K1 induced death15.8-fold faster than that of K28, killing 0.613% and 9.687% of the population per minute, respectively.

### 3.3. Integrated model of the primary pathway of toxic action

The integration of the binding models with the killing models for K1 and K28 necessitated the introduction of a single parameter, whose activity could be traced across both stages of binding followed by killing. In order to build the integrated model of toxin binding and toxin-induced death, the binding equations needed to be modified such that the binding was modelled based not on individual toxin molecules but on a cluster of toxin molecules represented by a single LU. In terms of the constants of the equation, this modification affected the saturation constants. Once a single LU was bound to the cell membrane, it was assumed to induce cell death subsequently, and the same LU was used up in the killing model to convert a viable cell into an inviable one. In this system both K1 and K28 were considered to be utilised only once, being bound on to the cell wall, and then on to the membrane, followed by its binding to the DNA. The toxin-induced cell death dynamics were represented by the following mass action kinetics in (eq. 7) and (eq. 8) for K and K28, respectively.

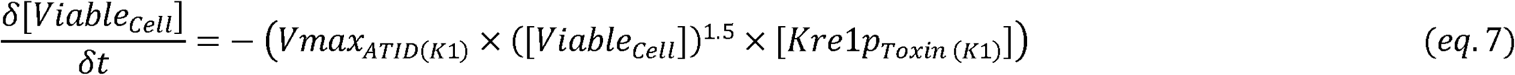

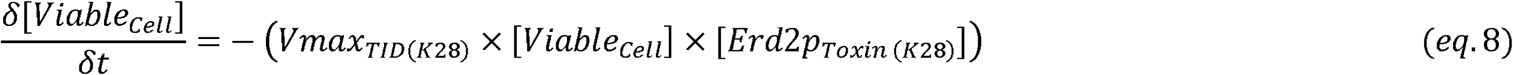

Kre1p and Erd2p are the respective cell membrane receptors of K1 and K28, and toxin bound receptor concentration represented the last step of the binding phase mechanism. [*Viable* _cell_] represented the size of the population of sensitive cells. [Kre1p_Toxin (Kl)_] and [Erd2p_Toxin (K28)_] represented the amount of K1 and K28 molecules bound to their respective receptors. The maximum specific rate of toxin-induced death,*Vmax*_*T ID*_, was determined by parameter optimisation.

The models of both K1 and K28 toxins successfully simulated toxin induced death dynamics reported earlier by [37] and [50] (Fig. 3a, 3b). The initial dynamics observed in the empirical data indicated faster cell death when the number of viable cells was higher for K1. Possibly, the high cell numbers could specifically facilitate the binding of an LU-equivalent number of toxin molecules to Kre1p with reduced competition. As the population of viable cells decreased, the ability for unbound K1 molecules to find a viable cell to bind to decreased accordingly. In order to achieve a model, which could represent this phenomenon, the order of the rate equation with respect to the viable cell concentration was set as 1.5 (eq.7), allowing rapid K1 killer activity from toxin binding on the cell wall and the cell membrane to the cell death. This modification was not needed for the case of K28, indicating K1’s superior efficiency in toxic activity, particularly during the initial period of the population’s exposure to toxins.

**Fig. 3.**
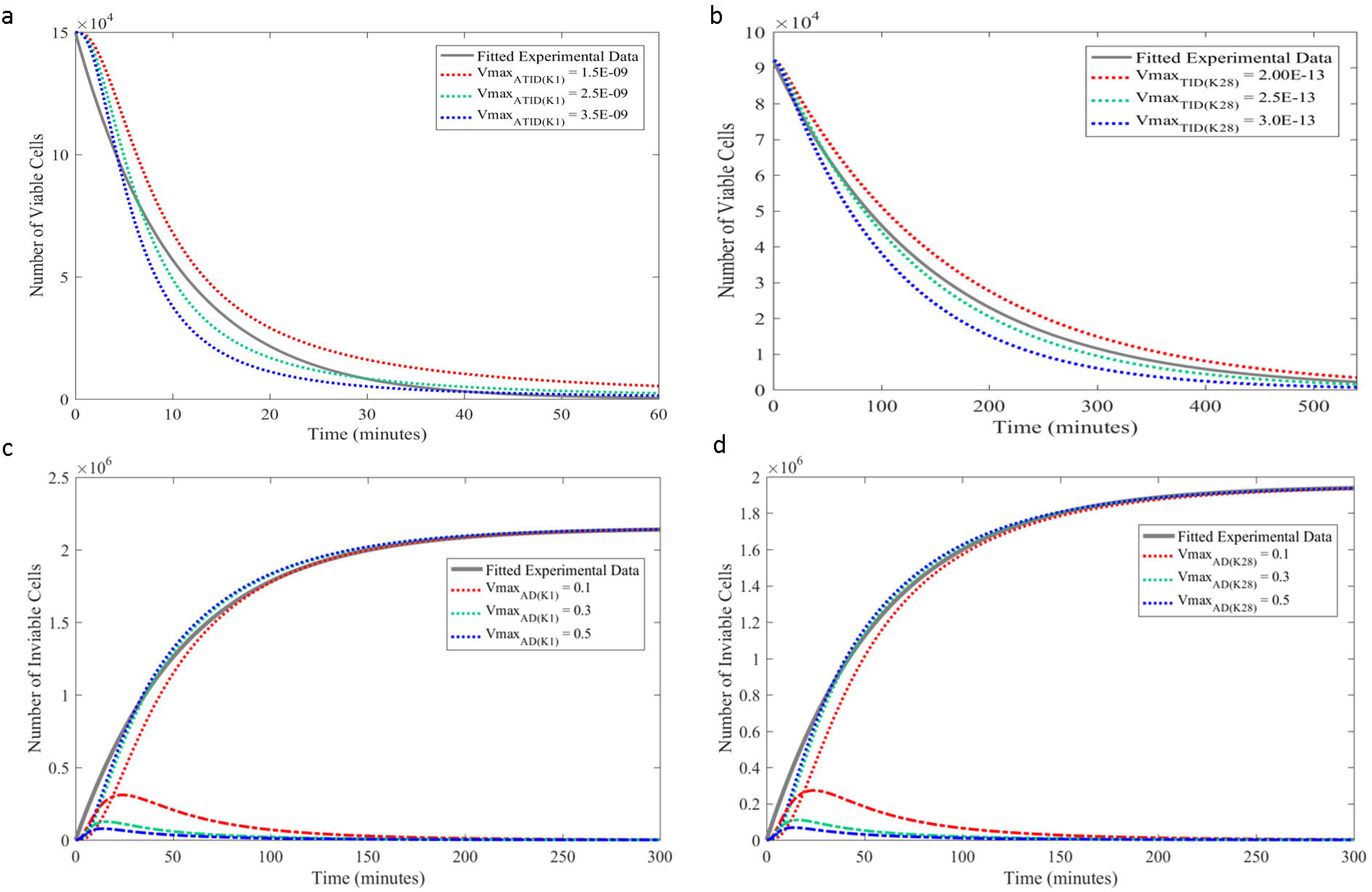
Modelling of the primary and secondary mechanisms of toxin-induced death. (a-b) Kinetic models of toxin-induced cell death via the primary action pathway for K1 (a) and K28 (b). The simulations were carried out utilising different maximum specific rate of cell death constants. The experimental data displayed in grey solid line was adopted from [37,50]. The model employed (eq.7) for (a) and (eq. 8) for (b). The total (bound + unbound) K1 concentration is 75000 toxin molecules/cell, and the total K28 concentration was 9.63×10□ toxin molecules/cell. (c-d) Kinetic models of toxin-induced cell death via the apoptotic action pathway for K1 (c) and K28 (d). The experimental data displayed in grey solid line was adopted from [37]. The model employed (eq.9, 10) for (c) and (eq. 11, 12) for (d). The dotted lines indicated the simulations concerning the inviable population, whereas the dashed lines denoted the fraction of the population that entered the apoptotic phase, but did not yet emerge as dead cells in (c) and (d). The toxin concentration necessary to induce death in a single cell is 2600 molecules in (c), and 2800 molecules in (d). All simulations were carried out utilising different maximum specific rate of cell death constants, and the simulations that used increasing values are shown in red to green to blue dashed lines in (a-d).

### 3.4. Integrated model of the apoptotic pathway of toxic action

Programmed cell death upon exposure to low concentration of toxins could be deemed a two stage process where an LU equivalent of toxin molecules bind to the cell membrane receptors triggering apoptosis, followed by cell death. Although the fate of the cell has already been determined at the initial stage, apoptosis requires a set of controlled actions to take place, introducing a time delay before the cells consigned to death can actually be considered as inviable. We presented this phenomenon in the model by introducing an additional model step to mark when cells would be considered as apoptotic. It represents the first stage of apoptosis described above, where the cells start showing signs of programmed cell death, and therefore are consigned to die, but are not technically dead yet. Toxins were modelled to induce cell death following a mass action rate law when at least a single LU equivalent of toxin modelcuels were bound to the cell membrane receptors, as in modelling of the primary toxin action pathway. The fate determination and death for K1 and K28 binding were represented by (eq. 9, 10) and (eq. 11, 12), respectively.

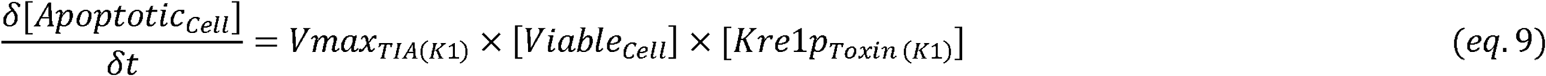

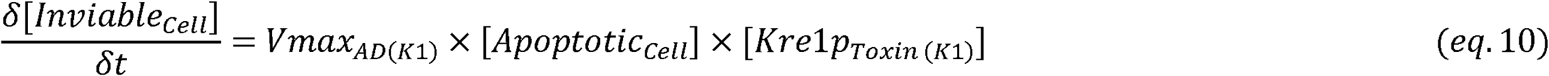

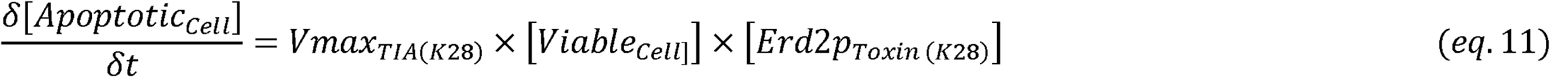

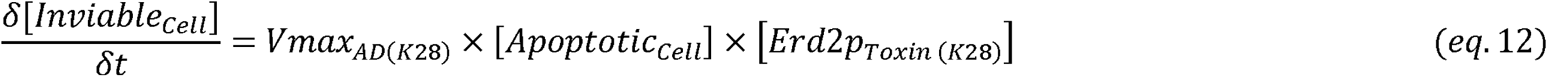

*Vmax*_*TIA*_ was determined *via* parameter estimation from [11] by identifying the duration for a cell that was exposed to 2800 PMKT2 molecules to displayed recognised apoptotic signals. The loss of viability was predicted successfully for both K1 and K28 (Fig. 3c, 3d). The initial killer activity of the model suggested a lag period before toxin-induced death was observed, which was likely to be missed in experiments due to incompatibility of the timescales. Experimental investigations to date focused on monitoring cell death over long time scales, and have failed to sample the population within the first couple of minutes upon toxin exposure. The model simulations were able to capture the delayed response in the early dynamics of the response. The transition from the apoptotic state to cell death was predicted to take place rather quickly by the model, as shown by the dynamic profile of the size of this intermediary population (Fig. 3c, 3d). Furthermore, a large variation in the maximum specific rate of death did not have a substantial impact on the dynamics of toxin-induced death, suggesting that the rate at which apoptosis was triggered in the sensitive cell population was the important step in this process. This further implied limited heterogeneity within the population as to how long apoptotic processes would take to kill the cell.

### 3.5. Modelling the killer activity of K1 and K28 toxins on sensitive populations of the same species

Generalised models of the killer toxin activity on sensitive yeasts were developed for two well-known killer toxins virally acquired by yeasts with distinct mechanisms of action; K1 and K28. Different stages and relevant design concerns discussed in Sections above were used to reconstruct a modular global model, and were then specialised for K1 and K28 activity accordingly. The different constants optimised and estimated for K1 and K28 from empirical data, which utilised in their respective models, are presented in Table 3.

The different modules were brought together under two conservation laws, leading to the final working models for simulating K1 and K28 activity: The total number of toxin molecules bound and unbound to receptors remained constant throughout the simulation, and the sensitive yeast population maintained steady growth, with natural death of the cells to replace the newly formed daughter cells, thus keeping the mean number and the mean size of the cells as well as the mean number of receptor molecules on cell wall and on the cell membrane constant unless exposed to killer toxins. These models were created and simulated using the open-source COPASI software, and the constructed models were made available both as Electronic Supplementary Materials 1 and 2. They were also deposited in the BioModels database with the following identifiers; MODEL1804230001 and MODEL1804230002 for K1 and K28, respectively. We also provide here a visual summary of the mathematical representation of both models separately, as these models possess a more complex universal structure than a simple compilation of the rate equations presented above (Fig.4).

**Fig. 4.**
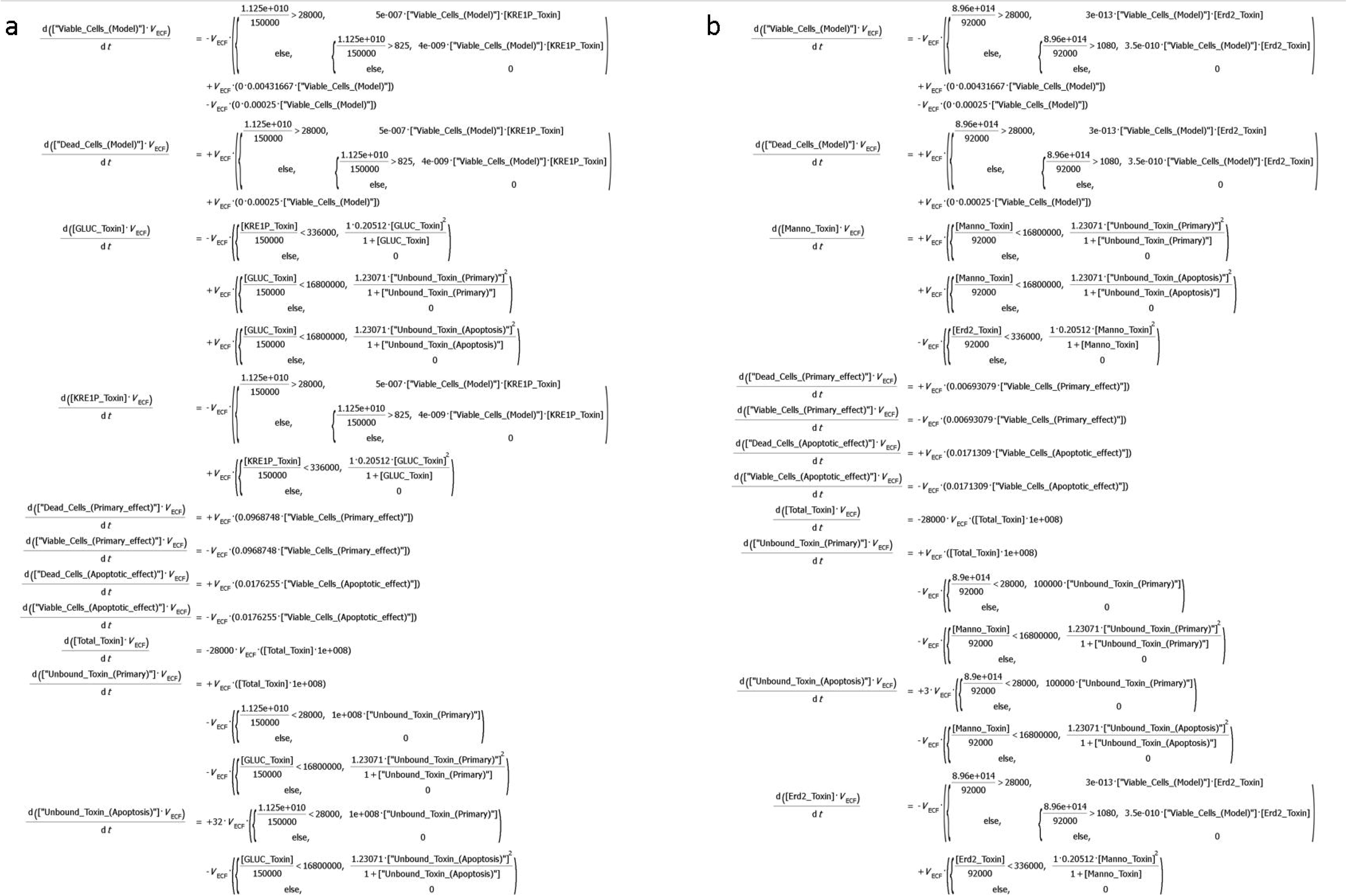
Summary of the mathematical representation of both models. A snapshot of the final model equations as implemented in COPASI are presented above. The set of equations in (a) and (b) relate to K1and K28 toxins, respectively. The editable version of these models can be accessed via COPASI software, by accessing the relevant models for each toxin.

## Discussion

We constructed a deterministic model, which predicted the killing dynamics of K1and K28 toxins, which agrees with the existing empirical data. Since the dynamics of K1 and K28 toxic activity was not investigated comparatively before, our model provided a useful platform to conduct a comparative analysis of the kinetics of these toxins, and their efficacy. The fastest toxic activity was observed in response to K1availability at high concentrations, when the primary toxin pathway was employed, and this was 5.5 times faster than its action when the apoptotic pathway was induced. Conversely, the route of action through the K28 primary pathway was approximately 3 times slower than the route of action *via* its apoptotic pathway. K1 available at high concentrations was more effective in killing the sensitive cell population than K28, when the primary pathways of toxicity were in order. However, the kinetics of the apoptotic pathways were essentially equivalent for both K1and K28. The large variability in the K28 primary pathway could be due to the fact K28 arrests cells specifically at the G1/S boundary phase of cell replication [11]. The rate at which cells enter the G1/S boundary phase would depend on environmental conditions and the state of the cell culture [52]. Therefore, varying experimental conditions across different datasets exploring K28 toxic activity would have led to these differing results.

The binding dynamics of the model showed that as long as the bound toxin concentration did saturate the surface receptors, the rate at which molecules adsorbed on the cell surface was independent of toxin concentration. Model predictions showed that the binding kinetics of the K28 primary pathway and that of PMKT2, the analogous toxin produced by *P. membranifaciens* were similar despite large differences in the available concentration of the two toxins, indicating that toxin-induced death was governed by the binding of a single LU-equivalent of toxin molecules, and that further increasing the amount of toxin bound did not necessarily speed up the rate of the process.

The empirical data on toxin binding, however, was largely disparate across different reports. Kurzweilová *et al*. reported that 16,800,000 molecules of K1 could bind to a single cell’s wall [25]. Recently, a modest figure of 555 K1 molecules/cell was proposed [27]. The number of β-1,6-D-glucans on a haploid parent yeast cell wall was estimated to vary in the range of 6,600,000 to 11,000,000 [26,53], and would be even higher for diploid cells than these reported here. Considering that a single toxin molecule can bind to any of these primary receptors, which are available most of the time, binding data reported by Kurzweilová *et al*. provided better representation of the binding kinetics for K1 than more recent reports did in the model simulations, which complemented the other empirical data, particularly on the mechanism of action of the apoptotic pathway.

K1 toxin was long disputed also to interact with the outward-rectifier potassium channel of the plasma membrane, Tok1p in addition to the Kre1p membrane receptor [54]. A counterargument was proposed by Breinig *et al*. proposing that Tok1p channels were only activated downstream, once K1 had already triggered ionophoric disruption [29]. The binding dynamics proposed by the model support the hypothesis that the Tok1p channels would be activated downstream along the toxin-induced death pathway rather than at the initial binding stage.

Several aspects of design were important for modelling the killer toxin dynamics. The total number of toxin molecules available, both bound to a receptor and unbound, was assumed to be constant according to the main conservation law. This conservation was built on two assumptions: The half-life of the killer toxins exceeded that of the simulation period, so that the effect of toxin protein degradation could be excluded. Although the authors are not aware of availability of such data for *S. cerevisiae* K1, K2 or K28, a study on a *Schwanniomyces occidentalis* killer protein, which displayed 75% identity and 83% similarity with killer toxin K2, was reported to have a half-life of at least 8 hours at 30°C at pH4.4 [55]; at a sufficiently close environment to those of yeast cultivations. The second assumption was on the killer yeast population being maintained at a steady rate, thus not introducing excess toxic proteins into the environment, as was discussed earlier. Furthermore, the toxin export through efflux pumps was excluded from the model, since there is no evidence yet in support of potential resistance mechanisms to be active against killer toxins.

Another important feature considered at the integration stage was the possibility of the recycling of toxin molecules. The notion of being able to reutilise toxin molecules, possibly similar to what the cell does in the case of the currency metabolites, was evaluated as a potential strategy. However, once passed through the nuclear membrane, K1 and K28 were reported to bind the DNA irreversibly [56], also undergoing an irreversible structural modification. Furthermore, no reports existed on the possibility of any reuse of toxins, and this notion was thus excluded from the design. Density of the cell culture, nutrient availability and dispersal were all shown to affect the competitive ability of toxin-producing yeasts [57,58], therefore, these parameters would potentially need to be taken into consideration for the specific applications.

A distinct threshold of the initial concentration of toxin molecules available per cell was adopted in this model to determine whether the toxic effect would be exerted *via* the primary or apoptotic pathway. Although this threshold was very useful for comparing and contrasting the kinetics of these two pathways of toxic activity, and represented the real kinetics of these mechanisms sufficiently well in light of existing empirical data, it should also be noted that simultaneous activity of the primary and the apoptotic pathways was also reported across a range of toxin concentrations; K1 was shown to kill cells *via* both the primary and apoptotic pathway in a range of concentrations varying from 28000 to 3000 toxin molecules per cell, but a “breaking point” between the two mechanisms was also reported in the same work at around 7500 molecules per cell where the pathway rapidly switched from one pathway to the other [37]. Although not of immediate concern due to lack of sufficient empirical data, the flexibility of the constructed models would allow the relevant modifications to be made to tailor the time and concentration dependence of killer toxin activity, if further experimental evidence backed up the transitional nature of these two mechanisms of action within proposed ranges of concentration profiles.

A stable killer phenotype has been known to necessitate a coordinated action by a group of host chromosomal genes including SEC genes, required for general secretion of extracellular proteins and glycoproteins, the KEX-encoded proteases for preprototoxin processing and precursor maturation of the yeast pheromone α-factor, in addition to a set of chromosomal genes, which either directly or indirectly affect dsRNA virus propagation, classified into two major groups; the maintenance of killer genes, MAK, and the superkiller genes, SKI [59]. The detailed mechanisms of action pertaining to these are beyond the scope of the work here, and were thus excluded from the models constructed in this work.

The initial phase of toxin induced death simulated by the model was observed to be consistently different from empirical observations, where a short lag phase was observed in model simulations introduced due to the binding of toxins prior to cell death, which was not observed experimentally. This discrepancy was thought to be caused by either of the two following reasons, or the combination thereof: (i) The deterministic nature of the model, which assumed synchronous action for all members of the cell population, failed to represent the heterogeneity caused by spatial or stochastic variations in actual systems. (ii) The most commonly employed experimental technique used in measuring cell death kinetics involves taking aliquots from the cell culture to measure the toxin activity at specific time points. It is quite common not to sample the culture within the first 15 minutes upon exposure, and therefore the experimental data available could just be an artefact of extrapolation as the true kinetics were not recorded during this time period. The model predictions, in fact, provide the missing details on the initial kinetics of cell death.

The action of cell membrane binding of K28 is complex as the toxin doesn’t simply bind to the cell periphery and trigger downstream effects. Instead, the toxin is endocytosed into the cell’s cytoplasm where it undergoes extensive modification before it can exert lethality in the cell’s nucleus. The mechanism of the pathway and most of the intracellular modifications taking place are now well-understood, albeit without any kinetic information available [18]. Therefore, all post-binding processes were lumped into the equations representing toxin-induced death in the current version of the model. Availability of kinetic data on this mechanism would allow the reconstruction and implementation of a more detailed model describing the physiology of toxin binding and cell death. The model of binding proposed in this work was nevertheless able to predict the overall killer activity of K28 successfully, however, it lacks sufficient details on the mechanistic action, which could only be implemented in light of future empirical kinetic data.

The models of killer toxin activity constructed here are able to simulate how the dynamics of sensitive yeast populations are affected by exposure to different toxins at various concentrations, enabling us to simulate excess environments as well as mildly toxic environments, which would be more likely to be encountered in the wild. These models (BioModels Database [42], MODEL1804230001 for K1and MODEL1804230002 for K28) constitute a useful platform to explore the dynamics of toxic activity and to offer insights to the mechanisms and suitability of each toxin and pathway for managing starter cultures. The predictions achieved from such models could be used to predict optimal ratios of killer and sensitive yeast cells, and propose control actions to maintain these optimal ratios. Accurate quantitative frameworks achieved through such models may assist building resilient and productive starter cultures of mixed fungal and yeast species for different biotechnological applications.

## Acknowledgements

The authors gratefully acknowledge the funding from the Leverhulme Trust (ECF-2016-681 to DD). The funding agency had no role in study design, data collection or interpretation, or the decision to submit the work for publication.

## Author contributions

DD designed and supervised the work. SS performed the modelling analysis. SS and DD drafted the manuscript. Both authors read and approved the final manuscript.

## Conflict of interest

The authors declare that they have no conflict of interest.

## Ethical approval

This article does not contain any studies with human participants or animals performed by any of the authors.

## Informed consent

All authors read and approved the manuscript.

## Electronic Supplementary Materials

### ESM1

Model of K1 killer activity

(.cps format, COPASI working file)

### ESM2

Model of K28 killer activity

(.cps format, COPASI working file)

